# Differential functional roles of fibroblasts and pericytes in the formation of tissue-engineered microvascular networks *in vitro*

**DOI:** 10.1101/558841

**Authors:** Natalia Kosyakova, Derek D. Kao, Francesc López-Giráldez, Susann Spindler, Morven Graham, Kevin J. James, Jee Won Shin, Xinran Liu, Gregory T. Tietjen, Jordan S. Pober, William G. Chang

## Abstract

**Aims:** Formation of a perfusable microvascular network (μVN) is critical for tissue engineering of solid organs. Stromal cells can support endothelial cell (EC) self-assembly into a μVN, but distinct stromal cell populations may play different roles in this process. Here we investigated the effects that two widely used stromal cells populations, fibroblasts (FBs) and pericytes (PCs), have on μVN formation.

**Methods and results:** We examined the effects of adding defined stromal cell populations on the self-assembly of ECs derived from human endothelial colony forming cells (ECFCs) into perfusable μVNs in fibrin gels cast within a microfluidics chamber. ECs alone fail to fully assemble a perfusable μVN. Human lung FBs stimulate the formation of EC lined μVNs within microfluidic devices. RNA-seq analysis suggested that FBs produce high levels of hepatocyte growth factor (HGF), and addition of recombinant HGF improved μVN formation within devices. Human placental PCs could not substitute for FBs, but in the presence of FBs, PCs closely associated with ECs, formed a common basement membrane, extended microfilaments intercellularly, and reduced microvessel diameters.

**Conclusions:** Different stromal cell types provide different functions in microvessel assembly by ECs. FBs support μVN formation by providing paracrine growth factors whereas PCs directly interact with ECs to modify microvascular morphology.

**Statement of Contribution:** Natalia Kosyakova, Derek Kao, William G. Chang were primarily responsible for the conception, design, interpretation of experiments, and drafting of the manuscript. Francesc López-Giráldez carried out analysis of RNA-seq data. Susann Spindler and Gregory Tietjen assisted with microvessel analysis software. Morven Graham and Xinran Liu assisted with the electron microscopy. Kevin J. James and Jee Won Shin assisted with data collection. Jordan Pober assisted with a critical review of manuscript and experimental design.

## Introduction

Microvascular network (μVN) formation is critically important for tissue engineering of organs too thick to be maintained by diffusive nutrient transport alone. We and others have generated human EC-derived microvascular networks (μVNs) *in vivo* within gels implanted into immunodeficient mice.^1^ However, human ECs suspended in the same gels *in vitro* initially assemble into cords but fail to fully form a μVN as the cells typically die between 24 and 36 hours. To improve vascularization, in previous experiments, we have over-expressed Bcl-2 to reduce the apoptotic response of human umbilical vein endothelial cells (HUVEC) in collagen/fibronectin matrices.^2, 3^ More recently, we have utilized human ECs differentiated from human endothelial colony forming cell (ECFCs) that also form vessels *in vivo* and have much greater replicative life spans than HUVECs, an important advantage for tissue engineering.^4^ However, like HUVECs, untransduced ECFC fail to form stable μVNs *in vitro*. Although Bcl-2 overexpression does not seem to cause transformation or give rise to tumors *in vivo*, there is still concern about this approach in clinically implanted tissues.

Microvessels are normally surrounded by extracellular matrix, stromal FBs, and supporting PCs that are intimately associated with the endothelium and share a common basement membrane. FBs are believed to be the principal cells of stromal tissue with critical roles in synthesis of extracellular matrix. FBs have key roles in the development and morphogenesis of tissues and organs.^5^ In contrast, PCs are critical for vascular development and for stabilization of the microcirculation. They are thought to regulate vascular tone, permeability, and have immunological functions.^6^ Genetic or acquired deficiencies in PC coverage of endothelial-lined capillaries result in abnormal microvasculature characterized by increased microvessel diameter and increased permeability.^7-9^

Thus, important biological questions arise about the roles of stromal cell types such as FBs and PCs in successful microvascular tissue engineering. In previous studies, FBs have been shown to support EC sprouting and lumen formation after being seeded onto collagen coated dextran beads within 3D fibrin gels. Secretion of FB factors is thought to be important in this angiogenic response.^10, 11^ We have also observed that these PCs invest tissue-engineered human microvasculature when implanted *in vivo*.^12, 13^ The presence of human PCs led to mural coverage, decreased vessel size, and permeability in tissue-engineered microvessels.^12^

Therefore, we believe that host stromal and EC interactions are critically important for the formation of μVNs. This is supported by observations that non-transduced human ECs formed robust μVNs when co-implanted with human mesenchymal stem cells,^14^ human lung FBs,^15, 16^ or mouse 10T1/2 cells.^17^ Others have described that human lung FBs can support EC survival and μVN formation in fibrin gels within a microfluidics device.^8^,^9^ These concepts and previous experimental investigations prompted us to compare the differential functions of FBs and PCs on ECFC-derived microvascular networks in an *in vitro* microfluidics chamber containing cells suspended in fibrin hydrogels. Here we report that these two stromal cell types are indeed distinct and play very different roles in μVN formation.

## Methods

### Primary cells, fluorescent labeling, and protein quantification

ECFCs were cultured in EGM-2MV (Lonza, Walkersville, MD, USA) on gelatin (Sigma-Aldrich, St. Louis, MO, USA)-coated plates and isolated from discarded and de-identified human umbilical vein cord blood as “late outgrowth” cells, as previously described.^4, 18^ Human microvascular placental PCs were isolated from discarded and de-identified placentas as explant-outgrowth cells, also as previously described.^12^ Both procedures were designated as not human research by the Yale Human Investigation Committee. Human lung FBs were purchased from Lonza. Both FBs and PCs were serially cultured in Medium 199 (Gibco, Grand Island, NY, USA) plus 20% FBS, 2 mM L-glutamine, 100 U/ml penicillin, and 100 μg/ml streptomycin (all from Invitrogen, Carlsbad, CA, USA). All cells were used between subculture 2-15.

Where indicated, cells were transduced with lentivirus (rLV.EF1.mCherry-9 or rLV.EF1.AmCyan1-9) to induce expression of mCherry or AmCyan per the vendor’s recommended protocol (Vectalys, Toulouse, France). Multiplicity of Infection (MOI) for mCherry ECFC and AmCyan PCs and FBs were 7 and 50.

For quantification of HGF secretion, 5.0 × 10^5^ PCs and FBs were grown in a single well of a 24-well dish for 24 hours and enzyme-linked immunosorbent assay (ELISA) was performed per manufacture protocol (R&D Systems, Minneapolis, MN, USA).

### Microfluidic device set up

The microfluidic devices contain six ports for loading cells, matrix, and media (AIM Biotech, Singapore). The undersides of the devices have permeable laminates that facilitate gas exchange while μVNs are being cultured. Each device contains a central channel (10.5 mm long and 1.3 mm wide) and two flanking media channels that are 0.5 mm wide. Central channels were loaded with cells and 2 mg/mL fibrinogen after addition of bovine thrombin (2 U/ml) (Sigma). After polymerization, EGM-2MV media was loaded into top channels. To change the media, the top two media wells were filled with 70 μL and 50 μL of media, left to right respectively, and the bottom two media wells were filled with 30 μL of EGM-2MV. Media in the wells were changed twice a day for the first three days, and then once a day for the rest of the duration of the experiment. To test effects of HGF on μVN formation, recombinant HGF (rHGF) (R&D Systems) at indicated concentrations was added to the media loaded into the microfluidic devices.

### Microscopy

For quantification of vessel densities, an epifluorescence microscope (Leica DMI6000, Wetzlar, Germany) was used to image the devices. Multiple individual images were tiled together to form composite images of the entire gel channel. For confocal imaging, a Leica TCS SP-5 Confocal Microscope was used. For time-lapse video images, a Zeiss Axiovert 200M inverted fluorescence microscope with a Hamamatsu ORC-AG high-resolution camera and Volocity imaging software (PerkinElmer, Waltham, MA, USA) was used to capture flow of Sphero™ 2 μm high intensity fluorescent beads (Spherotech, Lake Forest, IL, USA). For association studies, association was defined as direct contact between an AmCyan FB or AmCyan PC and an mCherry labeled microvessel. Field of interests were visualized with confocal microscopy. Z-stacks of serial sections were generated to capture the entire interaction and depth of the vessel. The 3D reconstruction of the vessel was analyzed for association.

For transmission electron microscopy, PCs and FBs were pre-labeled by incubating cells with 50 μg/ml of Molday ION Rhodamine B (BioPal, Worcester, MA, USA) for 18 hours in a single well of a 6-well plate. After the incubation period and three washes, cells were loaded into devices and μVNs were established and perfused for seven days before being fixed in 2.5% gluteraldehyde in 0.1 M sodium cacodylate buffer (pH 7.4) for 1 hour at room temperature. The devices were then rinsed in cacodylate buffer through the microfluidic device perfusion ports. At 1-hour post fixation, 1% osmium tetroxide was added, followed by rinsing and en-bloc staining in 2% aqueous uranyl acetate for an additional hour. Samples were then rinsed and dehydrated in an ethanol series. LRWhite (Electron Microscopy Sciences, Hatfield, PA, USA) acrylic resin was used to infiltrate the sample overnight followed by 48-hour incubation at 60**°**C. LRWhite was chosen over conventional epon resins because of its low viscosity that allowed for better flow into the thin microfluidic channels. Hardened blocks were cut using a Leica UltraCut UC7. Next, 60 nm sections were collected on formvar/carbon coated copper slot grids and contrast stained using 2% uranyl acetate and lead citrate. Samples were viewed on FEI Biotwin TEM at 80 Kv. Images were captured on Morada CCD and iTEM (Olympus) software. For EM tomography, 250 nm thick sections were collected using FEI Tecnai TF20 at 200 Kv with 15 nm fiducial gold to aid in alignment. Data was collected with SerialEM on a FEI Eagle 4×4 CCD camera using tilt angles of –60 to 60 degrees and reconstructed using IMOD. The 3D model was constructed using 3dmod software package in IMOD, following general modeling protocol. The slices within tomogram were manually drawn and contoured to generate a precise 3D reconstruction of the imaging sections.^19, 20^

### Microvessel density

To calculate the microvessel density, a self-developed MATLAB code with graphical user interface (GUI) was employed. The GUI was used to set parameters to filter out objects that were not microvessels (such as single cells). A gray threshold was used to create binary images. Single cells (which are smaller and rounder) were filtered out by setting a threshold on the minimum number of pixels as well as eccentricity of connected regions. Other regions (such as debris) were removed by manually drawing a region or targeting small areas for deletion. The vessel density was calculated as the number of pixels above threshold divided by the image area selected. The MATLAB code used is included in Supplemental Materials (Supplemental MATLAB Code).

### Statistical analysis

Statistical analysis was performed with Prism 7.04 (GraphPad) using two-way ANOVA with post hoc Bonferroni corrections. Three devices were analyzed per condition. Graphs are presented as means with standard deviations. When only two groups were compared, non paired two-tailed student’s t-test was performed.

### RNA-seq analysis

Confluent FBs from three different donors for each cell type were grown in one well of a 12-well plate and total RNA was purified using the RNeasy Mini Kit (Qiagen) with an on-column DNase treatment. Prep and sequencing was performed as described previously for PC RNA-seq analysis.^21^ For purified total RNA collected from FB samples, the three strand-specific sequencing libraries were produced following the Illumina TruSeq stranded protocol. According to Illumina protocol, the libraries underwent 76-bp paired-end sequencing using an Illumina HiSeq 2500, generating an average of 32 million paired-end reads per library. Both, the original PCs^19^ and the new FB sequences were processed together through the same analysis pipeline. For each read, the first 6 and the last nucleotides were trimmed to the point where the Phred score of an examined base fell below 20 using in-house scripts. If, after trimming, the read was shorter than 45 bp, the whole read was discarded. Trimmed reads were mapped to the human reference genome (hg38) with HISAT2 v2.1.0^22^ indicating that reads correspond to the reverse complement of the transcripts and reporting alignments tailored for transcript assemblers. Alignments with quality score below 20 were excluded from further analysis. Gene counts were produced with StringTie v1.3.3b^23^ and the Python script “prepDE.py” provided in the package. StringTie was limited to assemble reads matching the reference annotation GENCODE v27.^24^ After obtaining the matrix of read counts, differential expression analysis was conducted and normalized counts were produced using DESeq2.^25^ *P*-values were adjusted for multiple testing using the Benjamini-Hochberg procedure.^26^ Sequencing data for the FB samples were deposited in NCBI’s Gene Expression Omnibus (GEO) under accession number GSE122389.

## Results

### Co-culture of ECs and FBs within microfluidic devices

In previous studies, we have observed that implantation of ECFCs alone *in vivo* leads to formation of robust μVNs.^4, 27^ However, within microfluidic devices, ECFCs alone are not sufficient to form μVNs (Fig. 1A). Indeed, ECFC cords started to emerge by day 3, but began to deteriorate by day 5 and never achieved perfusable networks. Next, we investigated whether addition of a commonly utilized, commercially available normal human lung FBs could improve μVN formation. We found that a minimum of 2.5 × 10^5^ FBs per ml were needed to support the formation of stable perfusable μVNs. Increasing FB concentration up to 2.5 × 10^6^ FBs per ml improved microvessel density (Fig.1B). In subsequent experiments, we used 2.5 × 10^6^ FBs per ml of matrix when co-cultured with ECFCs. We confirmed that μVNs were perfusable in the ECFC and FB co-cultured devices by flowing fluorescently labeled beads through the channels (Supplemental Video 1). Although most experiments we performed were in cultured devices for 7 days, we have observed that co-culture of ECFCs with FBs supports the formation of perfusable μVNs that were maintained for up to 27 days in fibrin gels, but demonstrated declining microvascular density after day 9 (Fig. 1C).

**Figure 1.**
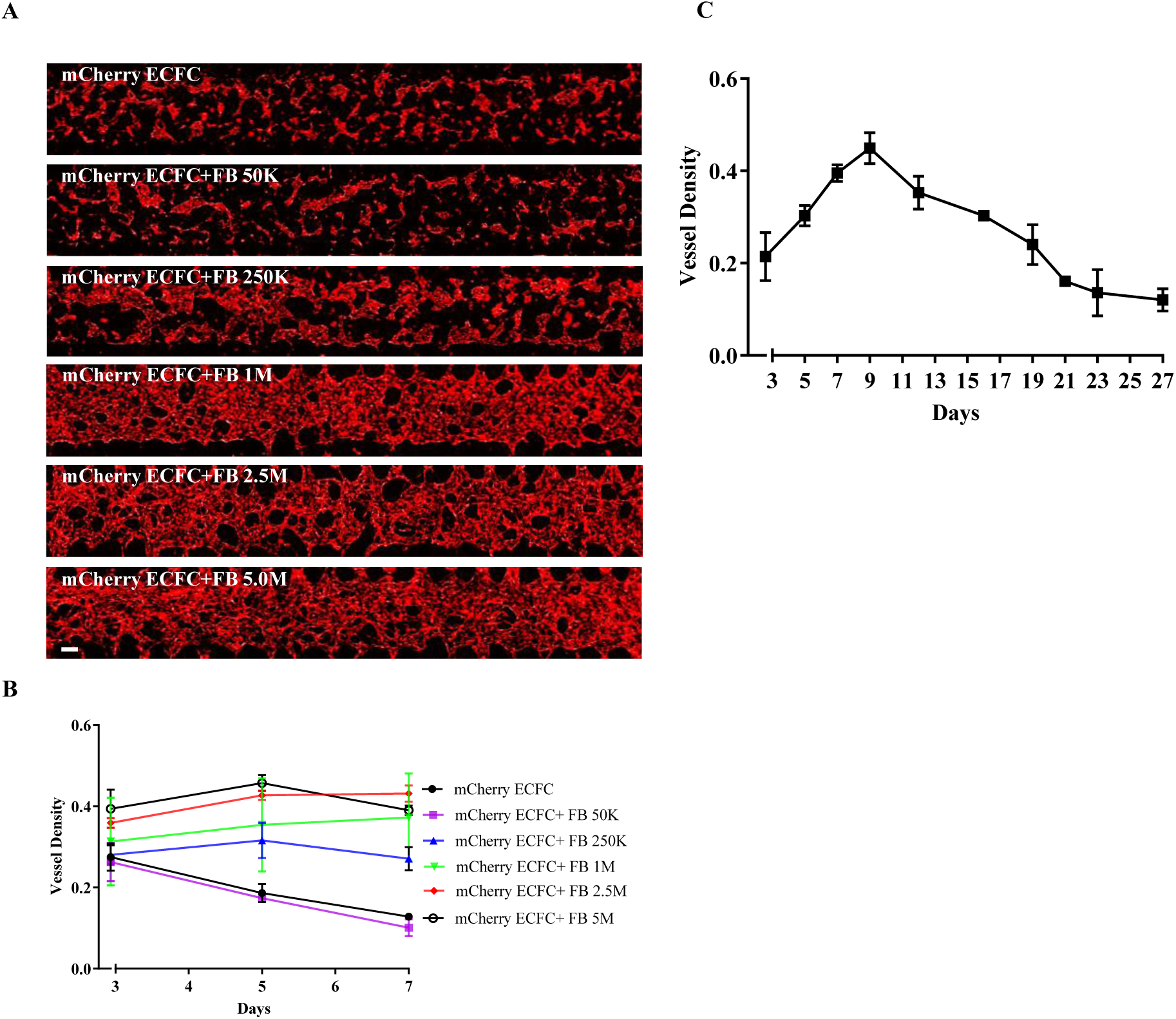
FBs stabilize μVNs. (A) Microfluidic devices with FB titration cultured for 7 days. ECFCs were 1.0 × 10^7^ cells per ml and FBs ranged from 5.0 × 10^5^ to 5.0 × 10^6^. (B) Vessel density within microfluidic devices. Statistically significant difference exists between ECFC only sample and samples containing2.5 × 10^5^ FBs per ml at Day 5 and 7 (*p*-value<0.0001). (C) Long term culture of microfluidic devices with 1.0 × 10^7^ ECFCs per ml and 2.5 × 10^6^ per ml FBs. Scale bar is 250µm.

### Co-culture of PC modified microvessel diameters

Given that PCs are known to stabilize microvessels *in vivo*, we investigated whether PCs co-cultured with ECFCs could also support the formation and maintenance of microvessels. Surprisingly, unlike FBs, PCs co-cultured with ECFCs did not yield stable μVNs (Fig 2A).

**Figure 2.**
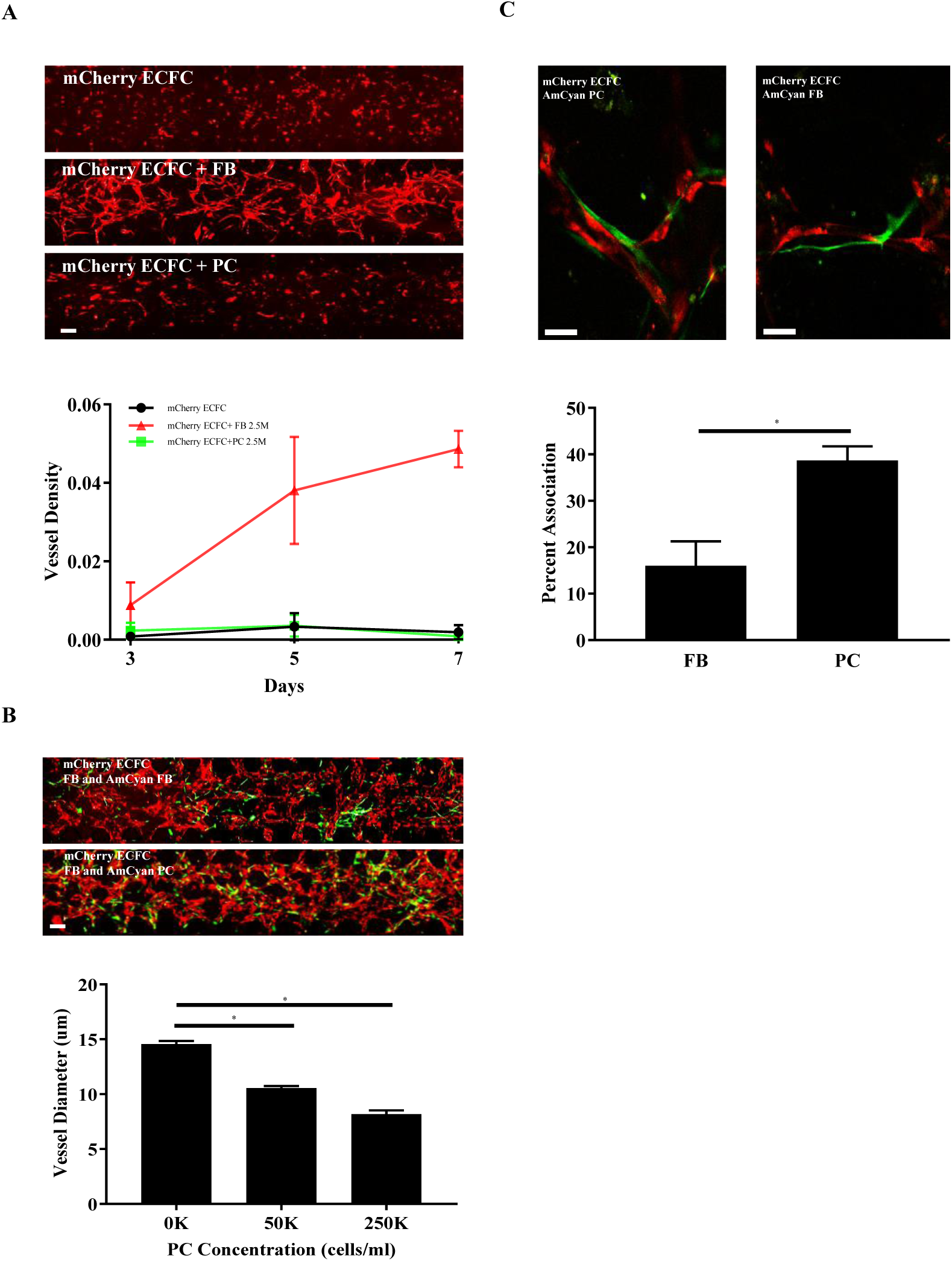
PCs are closely associated with and reduce the lumen diameters of μVNs. Microfluidic devices cultured for 7 days with 1.0 × 10^7^ ECFCs per ml alone and ECFCs with 2.5 × 10^6^ per ml PCs or FBs. Vessel density quantification of microfluidic devices completed with graphical user interface within MATLAB program. Statistically significant difference observed at Day 5 and 7 for ECFC+FB when compared to ECFC alone and ECFC+PC (*p*-value<0.0001). (B) Microfluidic devices cultured for 7 days with 1.0 × 10^7^ ECFCs per ml with 2.5 × 10^6^ per ml of FBs and 2.5 × 10^5^ of AmCyan FBs or 2.5 × 10^6^ per ml of FBs and 2.5 × 10^5^ AmCyan PCs. With 1.0 × 10^7^ ECFCs per ml and 2.5 × 10^6^ per ml FBs, increasing amounts of PCs reduce microvessel lumen diameters (*= *p*-value<0.0001). (C) PCs are significantly more associated with microvessels than FBs (*=*p*-value<0.003). Confocal images of ECFC (mCherry, red) with PC or Fib (AmCyan, green). Scale bars are 250µm (A, B) and 75µm (C).

Having observed that ECFCs and PCs were not sufficient to form stable μVNs, we co-cultured PCs with ECFCs and FBs to examine the effects of PCs on the μVNs (Fig. 2B). We observed that the addition of PCs reduced the diameters of the microvessels formed (Fig. 2B). The average diameter of the microvessels with no PC incorporation was 14.55 ± 0.283 μm, however the mean diameter of μVNs with 2.5 × 10^5^ PCs per ml were 8.18 ± 0.366 μm. In addition, there was an observable dose effect as diameters decreased with increasing PC number. We also observed that PCs were more closely associated with the microvessels formed than FBs when we examined μVNs by confocal microscopy (Fig. 2C). Fifty AmCyan fluorescently labeled FBs or PCs were examined by confocal microscopy and were identified as either associated with mCherry labeled microvessel or not associated. Of the counted cells, 16.00 ± 5.29% of FBs were associated with microvessels, while 38.67 ± 3.06% of PCs were associated with microvessels. PCs were significantly more likely to be associated with microvessels (Fig. 2C).

### RNA-seq comparison of FBs and PCs

Given the differences that we observed within the microfluidic devices when ECFCs were co-cultured with FBs or PCs, we used RNA-seq to compare bulk gene expression profiles of the FBs and PCs. We prepared and analyzed FB gene expression profiles with RNA-seq using three different donors in the same manner as previously published for PCs^21^ (Supplemental Table 1). We identified 1056 genes that were differentially expressed (*q*-value <0.05 and log^2^[fold change]≥ 5; Fig. 3A). Of the differentially expressed genes, we searched for proteins that may contributed to the phenotypic differences that we observed between FBs and PCs. In static endothelial sprouting models, others have identified factors that contribute both to vessel sprouting (angiopoietin-1, angiogenin, HGF, transforming growth factor-α, and tumor necrosis factor) and lumen formation (collagen I, procollagen C endopeptidase enhancer 1, secreted protein acidic and rich in cysteine [SPARC], transforming growth factor-β induced protein ig-h3 [βig-h3], and insulin growth-binding protein 7).^10^ When we specifically profiled the FB factors previously reported to enhance microvessel formation,^10^ we observed that HGF and βig-h3 or transforming growth factor β induced (TGFBI) as it is now named were significantly upregulated in FBs (Fig 3B). Collagen I expression was high in both FBs and PCs cultured as monolayers. Given the close association of PCs to ECFCs within the μVNs, we looked at subsets of genes involved in cell-cell, cell-matrix adhesion, and extracellular matrix proteins using gene ontology (GO0098609, GO0007160, and GO0031012). We observed that several integrin subunits, matrix adhesion genes, and basement membrane proteins were upregulated in PCs (Fig. 3C and Supplemental Table 2).

**Figure 3.**
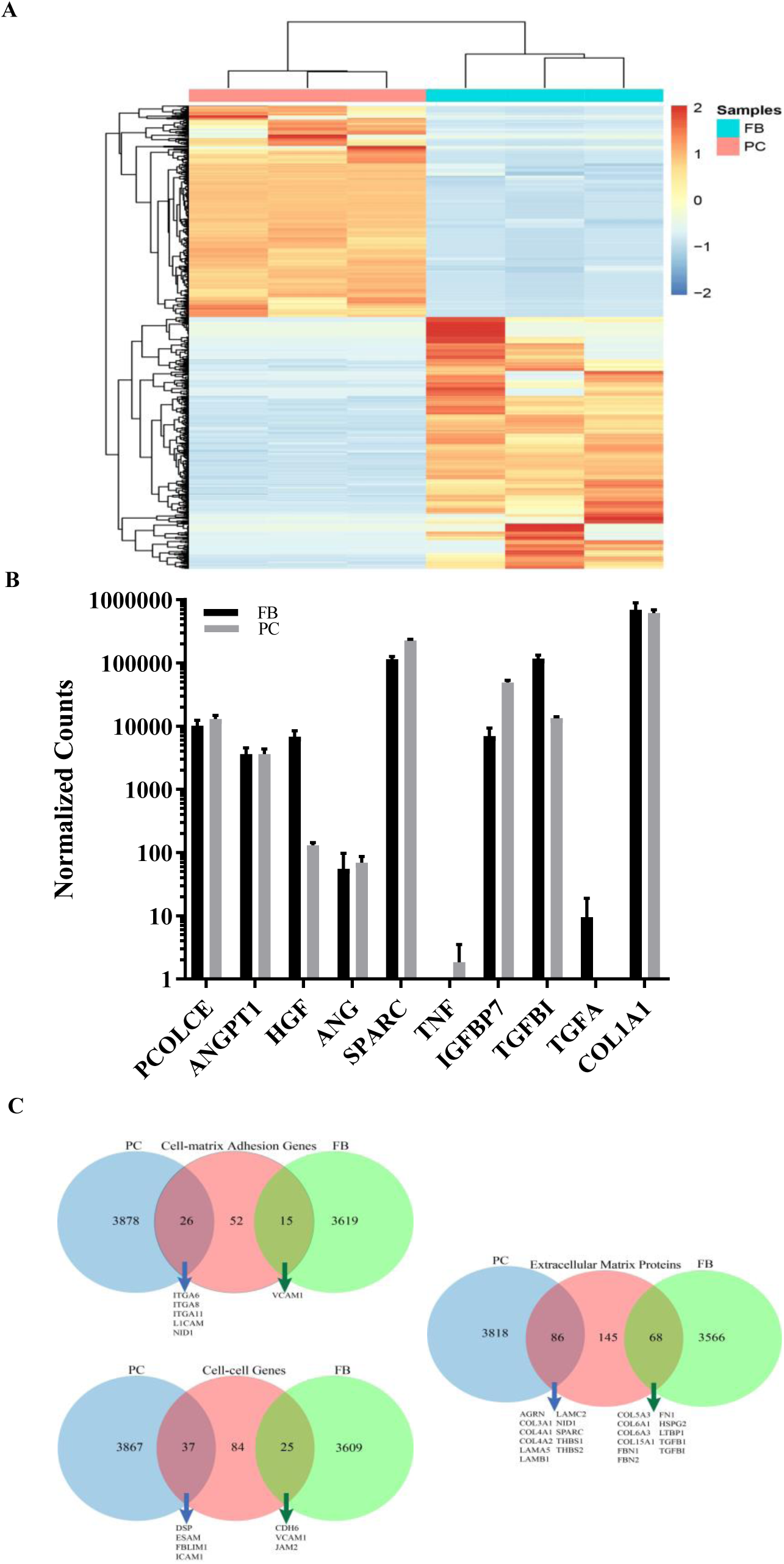
RNA-seq analysis of FBs versus PCs. (A) Heatmap of the gene expression profiles of FBs and PCs. Genes significantly differentially expressed (*q*-value <0.05) with a log^2^[fold change] ≥ 5 between FBs and PCs are shown (*n* = 1056). Cells from individual donors are represented in each column. Each row represents gene expressions across both cell types. Rows and columns are hierarchically clustered. Expression values are scaled by row; red color indicates higher gene expression and blue indicates lower gene expression. (B) Regularized log counts with standard deviations of FB factors previously shown to have positive effects on microvessel formation. (C) Venn diagrams show the number of genes differentially upregulated (*q*-value <0.05) in PC (blue) and FB (green) cell types. Red circles indicate number of genes which encode for subset of cell-cell, cell-matrix adhesion, and extracellular matrix proteins based on gene ontology (GO) terms. The intersecting regions of circles indicate the number of genes differentially upregulated in PC or FB within the GO subset. Genes of interest are highlighted.

### HGF stimulates μVNs in microfluidic devices

While a completed analysis of genes differentially expressed by FBs and PCs is beyond the scope of this current series of experiments, we did seek to establish whether the microfluidic devices could be used as a tool for dissecting how FBs contribute to stabilization of μVNs. Of FB factors reported to stimulate microvessel formation *in vitro*, HGF emerged as most significantly different. HGF has been demonstrated to stimulate blood vessel formation^28, 29^ and has important roles in development, cell survival, and tissue regeneration.^30^ *HGF* had an approximately 50-fold higher gene expression in FBs than PCs. We observed by ELISA that ∼5-fold higher levels of HGF protein was secreted by FBs than PCs (Fig. 4A). When rHGF was added to the culture media of the microfluidic devices, we observed an improvement in μVN density (Fig. 4B). However, even at high doses, we were not able to achieve the density and sustainability of μVNs formed by the co-culture of FBs (Fig. 4C). This suggests that FBs contribute other factors important to μVNs formation and stability.

**Figure 4.**
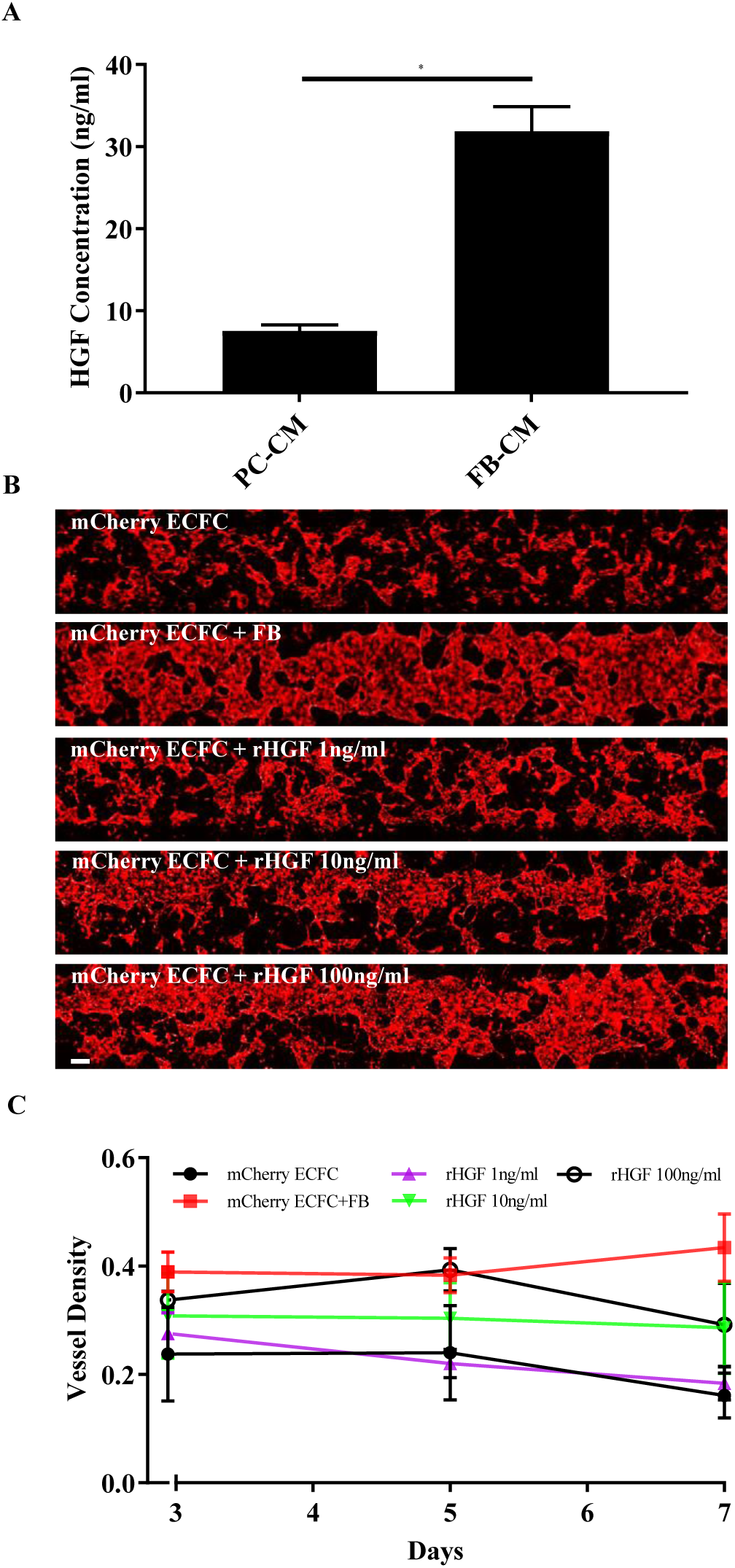
HGF stimulates μVNs. (A) FBs secrete more HGF as measured by ELISA quantification (*p*-value<0.0002). (B) Microfluidic devices when rHGF is added to the media (at indicated concentrations) compared to ECFC alone and ECFC with co-cultured FBs. (C) Vessel density improves with the addition of rHGF. Statistically significant difference exists between ECFC only condition and that containing 100 ng/ml of rHGF at Day 5 and 7 (*p*-value<0.05). Scale bar is 250µm.

### Ultrastructural analysis of μVNs

To further investigate the differences between PC and FB interactions within microvascular networks, we performed transmission electron microscopy (TEM) on the μVNs within the microfluidic devices at the 7-day time point. To distinguish the different cell types, we pre-labeled FBs and PCs with Molday ION (iron oxide particles). These particles accumulate in endosomes and are readily detected in TEM as electron dense particles within endosomes. With TEM, we rarely observed occurrences where the ECs and FBs were closely associated (Fig. 5A). When we did see associations, the cells remained distinct with the presence of collagen fibers between them. However, there was a much more dynamic interaction between ECs and PCs with formation of microfilaments between the two. In some cases, both cell types appeared to protrude actin filaments towards each other. In addition, we observed instances where a more established common basement membrane had formed (Fig. 5B and 5C). To better analyze the interaction between ECs and PCs, we used electron tomography to obtain a 3D delineation of the boundary between these cells (Fig. 6 and Supplemental Video 2). We observed that both cell types appeared to extend microfilaments towards each other, some were shared by the two cells and in specific places, cells were able to contact each other.

**Figure 5.**
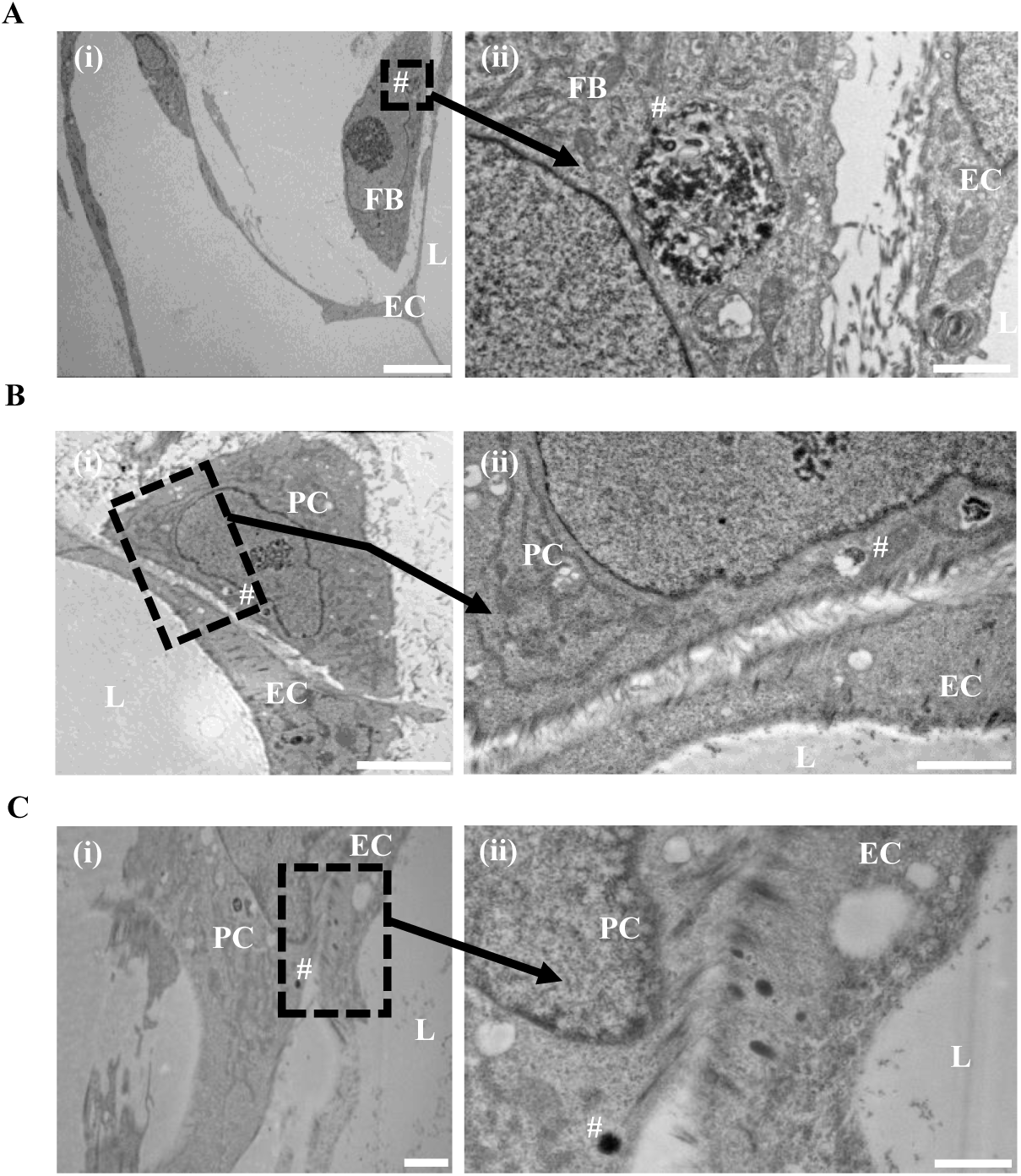
Ultrastructural analysis of cell-cell interactions within μVNs. PCs and FBs were pre-labeled with Molday ION particles prior to formation of μVNs and analyzed by TEM. (A) Association between ECFC and FB. Inset shows iron labeled endosome within FB and collagen fibers between ECFC and FB. (B) Interaction between ECFC and PC. Inset shows dynamic interaction observed between ECFC and PC. (C) Iron-labeled PC with ECFC. Inset shows established common basement between two interacting cells. Scale bars indicate 10µm (A (i), B(i)), 2µm (B (ii), C (i)), and 1µm (A (ii), C (ii)). Symbols indicate EC = ECFC, L=lumen, and #= electron dense particles due to iron labeling.

**Figure 6.**
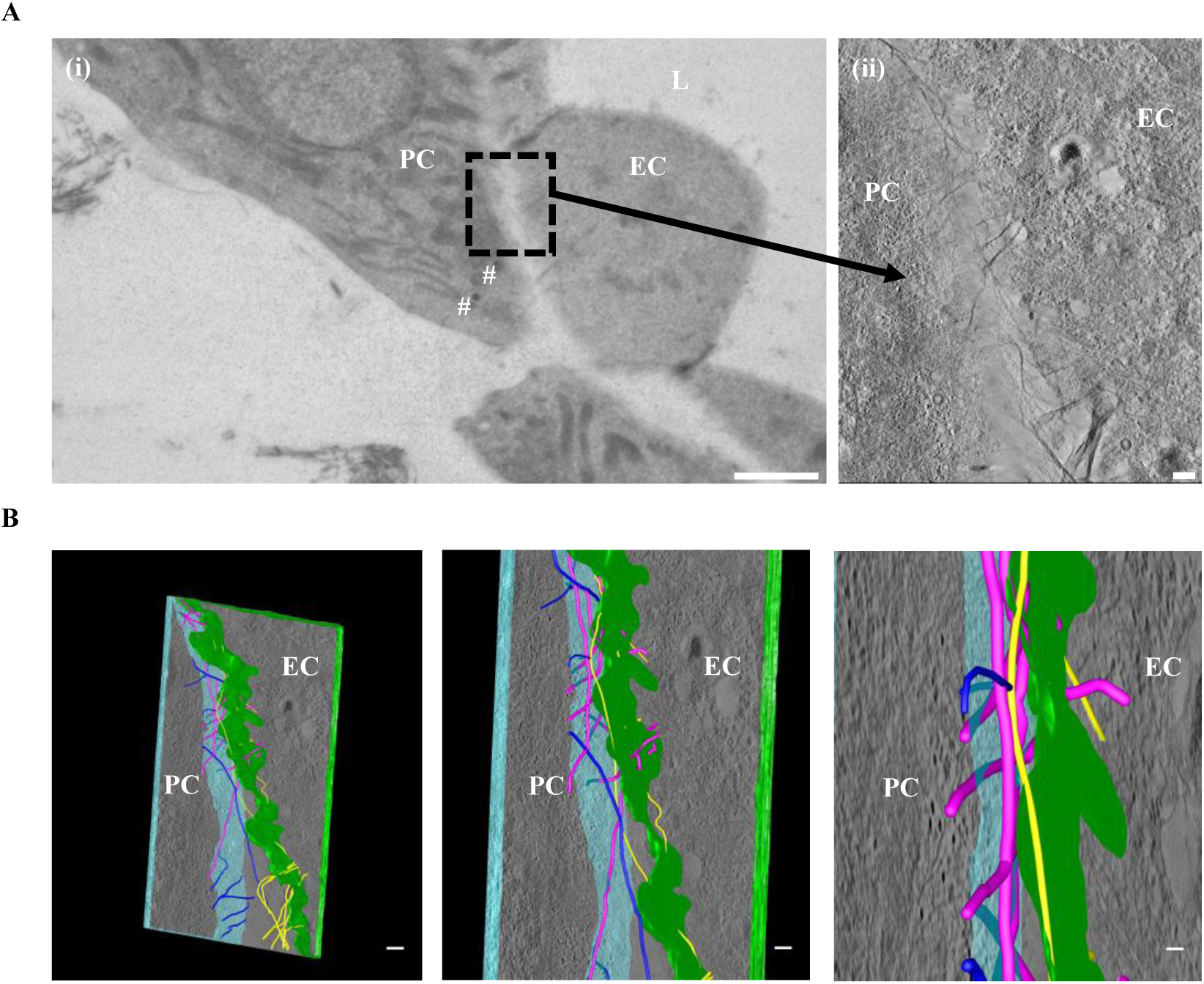
Electron tomography of ECFC and PC interaction. (A)Visualization of the field used to create tomography slices. Higher magnification area indicates the area which tomographic slices were created from and modeled using iMOD software (inset). (B)3D model reconstruction of tomography slices. ECFC border was outlined in green and PC in blue. Microfilaments are outlined in three colors: dark blue, magenta, and yellow to distinguish the location filaments originate from. Dark blue filaments originate in PC and yellow filaments originate in ECFC and do not cross over into neighboring cell. Magenta filaments connect both cell types. Symbols indicate EC= ECFC, L=lumen, and #= electron dense particles due to iron labeling. Scale 10um (A (i)), 200nm (A (ii), B).

## Discussion

A better understanding of how μVNs form is critical to advancing tissue engineering and future regenerative medicine approaches. However, the assembly of μVNs is a complex process with multiple cell-cell, cell-matrix, and signaling pathways necessary for success. Here we have examined how two different stromal cell types, FBs and PCs, affect formation of μVNs. We observed that FBs sustain and stimulate μVNs formation through paracrine release of survival factors, while PCs directly contact ECFCs to modify microvessel morphology.

A key issue raised in this study is the type of cells to use for tissue engineering research of μVNs. The same types of cells from different tissue sources may significantly vary in their behaviors. We chose human ECs derived from cord blood ECFCs, human lung FBs, and human placental PCs. We selected this EC population because it displays extended replicative lifespan and robust vasculogenic potential, is readily obtained, and can be genetically modified and clonally selected after CRISPR/Cas9 implementation.^4^ We selected human lung FBs because of their wide-spread use in experimental settings of tissue engineering. Finally, we have chosen placental PCs because they are also readily obtained from discarded post-partum tissue and have been characterized both behaviorally and by transcriptomics.^21^ We analyzed the interactions of these cells within gels cast in microfluidics chambers because this *in vitro* setting gives rise to perfusable μVNs, better recapitulating the *in vivo* setting.

The RNA-seq analyses demonstrate that FBs and PCs possess distinct gene expression patterns that likely contribute to the differences in their effects on μVNs within these devices. Looking at previously reported FB factors^10^ that improve microvessel formation, TGFBI and HGF was significantly upregulated in FBs compared to PCs. Furthermore, when rHGF was added to the μVNs without FBs, there was improvement in microvessel density. However, rHGF was not sufficient to fully reproduce the effect of FBs within the device, suggesting that there are other factors important for μVNs formation. TGBI is induced by TGF-β1 and β2 and is a secreted extracellular matrix (ECM) protein involved in morphogenesis, adhesion/migration, tumorigenesis, wound healing, and inflammation. Clinically, mutations of this gene lead to corneal dystrophy. Interestingly, TGFBI has been described to inhibit cellular adhesion to ECM^31^ and may be important for EC tubulogenesis.^32^ An alternative explanation for the inability of PCs to support μVNs formation in the absence of FBs is that the PC gene products may inhibit microvessel formation via cell-cell or cell-matrix interactions, yet still mediate modification of microvessel morphology. Several candidate proteins emerged from the RNA-seq data that could play a role in interactions between ECFCs and PCs. For instance, desmoplakin and several integrin subunits (*ITGA1, ITGA3, ITGA6, ITGA8, ITGAV, ITGB3, ITGB5*) were significantly higher in PCs and could be involved in ECFC-PC cell junctions or PC matrix interactions. More specifically, desmoplakin has been reported to interact with intermediate filaments and N-cadherin^33^ and is thought to be a key regulator of cell mechanics.^34^ Furthermore, N-cadherin is thought to be important for EC anchorage to PCs.^35, 36^ In addition, integrins are heterodimeric proteins critical for cell-matrix interactions, mechanotransduction, and cell signaling.^37^ Consistent with our observation that ECFC and PCs form a common basement membrane, several ECM proteins were upregulated in PCs that are known to be part of vascular basement membranes^38^ including collagen type IV isoforms, laminin chains, nidogen 1, SPARC, agrin, fibulin 2, and thrombospondin 1 and 2. In future studies, we will further investigate the critical factors that mediate the disparate interactions between ECFCs, FBs, and PCs.

The recent emergence of microfluidic technologies allows for improved control of complex cellular interactions and fluid flows at the micro-scale. Additionally, microfluidic devices have low costs, are readily visualized in real-time, and the matrix and cellular components within can be readily modified. The 3D orientation and perfusability of the microvessels formed within microfluidic devices allows investigators to observe cell-cell interactions and to test microvessel characteristics in ways that cannot be accomplished in traditional 2D monolayer or co-culture systems. Interactions between ECFCs and PCs are clearly distinct from ECFCs and FBs. Furthermore, our electron microscopy studies appear to show the dynamic interaction and the formation of a common basement membrane between PCs and ECFCs. These images are reminiscent of reported TEM images of ECFCs and PCs *in vivo*.^39^ To our knowledge this is the first report demonstrating this process at the ultrastructural level within microfluidic devices. The observation of microfilaments and protrusions from these cells towards each other suggests a dynamic interaction that will need to be investigated further. The techniques used offer important experimental systems and tools for a deeper understanding of the interactions between the different cell types required for assembly of μVNs.

In conclusion, the findings reported here can be used to better inform the design and implementation of tissue-engineered μVNs in the future. Specifically, both FBs and PCs are likely necessary for tissue engineering of physiological human μVNs because these stromal populations provide different functions. FBs support formation and survival of μVNs whereas PCs have an important structural role. Each cell type has a distinct phenotypic and transcriptional profile that argues that they are not interchangeable in microvascular tissue engineering.

## Funding

This work was supported by the National Institutes of Health [WGC by K08EB018978 and JSP by R01 HL085416]. JS was supported by the Yale Science and Engineering Association Undergraduate Research Grant.

## Acknowledgements

Sequencing was performed at the Yale Center for Genome Analysis (YCGA) facility. We thank the Yale Center for Research Computing for guidance and use of the research computing infrastructure.

## Conflict of Interest

None declared

## Data Availability

Representative images, analyses, code, and expression data are included in this published article (and its supplementary information files). Raw image files are available from the corresponding author on reasonable request.

**Supplemental MATLAB Code**. Microvessel analysis code.

**Supplemental Figure 1. Microvascular density quantification**. Tiled images of μVNs were uploaded into the MATLAB program. Graphical user interface (GUI) was used to set thresholds for vessel density calculations.

**Supplemental Video 1. Perfusion through μVNs.** High intensity fluorescent beads (2 μm) were perfused through μVNs to confirm perfusability.

**Supplemental Video 2. iMOD 3D Modeling of EC and PC interaction.** EC border (in green),PC border (in blue), and microfilaments between the cells were manually contoured and meshed to construct a 3D model. The model was constructed in 3dmod, an image processing dialog found under iMOD programs. The tomography slices were opened in 3dmod and the outline of the cells was modeled in the main 3dmod window as a closed object. The microfilaments were modeled using the slicer tool which allowed us to follow and adjust for the x, y, and z orientation of the filaments. The contours were then meshed together and viewed in the model window. To better distinguish the origin of the microfilaments located between cells, microfilaments were outlined in three colors: dark blue, yellow, and magenta. Dark blue and yellow filaments originate in PC and EC, respectively. Magenta filaments connect EC and PC. Scale bar is 200nm.

**Supplemental Table 1.** RNA-seq comparison of human placental PCs and human lung FBs.

**Supplemental Table 2.** List of upregulated cell-cell adhesion and cell-matrix genes, and extracellular matrix proteins in PC and FB cell types. List generated using gene ontology resource.

