## Supplemental Figures for "Differential functional roles of fibroblasts and pericytes in the formation of tissue-engineered microvascular networks *in vitro*"

### Supplemental MATLAB Code

```
clear all
close all

images = dir('*cut.tif');

% parameters found in GUI:
thresh       = 0.1; % for binarizing image
pixel_thresh = 500; % minimum number of pixels for counting a region as vessel
ecc_thresh   = 0.4; % minimum threshold to exclude round structures

mark_objects                = 1; % possibility to mark and remove objects manually
draw_boarder_around_objects = 1; % possibility to draw boarder around regions to exclude

for i = 1:length(images)

    img_name   = images(i).name;
    img        = imread(img_name);     % load in image (4 channels)
    img        = img(:,:,1);           % only first channel is important
    img_height = size(img,1);
    img_width  = size(img,2);

    figure(1)
    imshow(img)

    img_bw       = imbinarize(img,thresh); % binarize image
    img_cr       = bwconncomp(img_bw); % find connected regions
    numPixels    = cellfun(@numel,img_cr.PixelIdxList); % number of pixels for each connected region
    eccentricity = regionprops(img_cr,'Eccentricity'); % eccentricity for each connected region
    ecc          = struct2cell(eccentricity);

    % filter out objects which are not vessels based on their size and
    % eccentricity
    img_detected_vessels = zeros(img_height, img_width);
    for j = 1:img_cr.NumObjects
         if numPixels(j) > pixel_thresh && ecc{1,j} >= ecc_thresh
            img_detected_vessels(img_cr.PixelIdxList{j}) = 1;
         end
    end

    figure(2)
    imshow(img_detected_vessels)

    %remove isolated objects manually:
    if mark_objects == 1
       index = 1;
       h = msgbox('Mark region')
       [x,y] = ginput(1);
       m = round(y);
       n = round(x);
       linindex = sub2ind([img_height, img_width],m, n);
       positions_list(index) = linindex;

       choice = questdlg('Mark another region?','Message box', 'Yes', 'No','Yes')
       while strcmpi(choice,'Yes');
             index = index + 1;
             [x,y] = ginput(1);
             m = round(y);
             n = round(x);
             linindex = sub2ind([img_height, img_width],m, n);  % conversion of x,y coordinates into linear indices
             positions_list(index) = linindex;
             choice = questdlg('Mark another region?','Message box', 'Yes', 'No', 'Yes')
       end

       for k = 1:img_cr.NumObjects   % remove all marked regions
           exist = [];
           compare = ismember(linindex,img_cr.PixelIdxList{k});
           exist = find(compare);
           if length(exist)~= 0
               img_detected_vessels(img_cr.PixelIdxList{k}) = 0;
           end
       end
    end

    % remove objects connected to vessels by drawing a boarder:
    if draw_boarder_around_objects == 1
        index = 1;
        h = msgbox('Select region')
        selected_region{index} = roipoly();
        choice = questdlg('Select another region?','Message box', 'Yes', 'No','Yes')
        while strcmpi(choice,'Yes');
              index = index + 1;
              selected_region{index} = roipoly();
              choice = questdlg('Select another region?','Message box', 'Yes', 'No', 'Yes')
        end
        selected_regions = selected_region{1};
        for l = 2:length(selected_region)
            selected_regions = selected_regions + selected_region{l};
        end

        img_detected_vessels = img_detected_vessels-selected_regions;
    end

    figure(2)
    imshow(img_detected_vessels)

    vessel_density(i) = nnz(img_detected_vessels)/(img_height*img_width);
end
```

Supplemental Figure 01

**A**

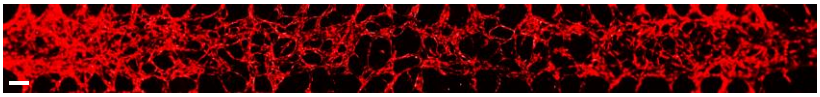

**B**

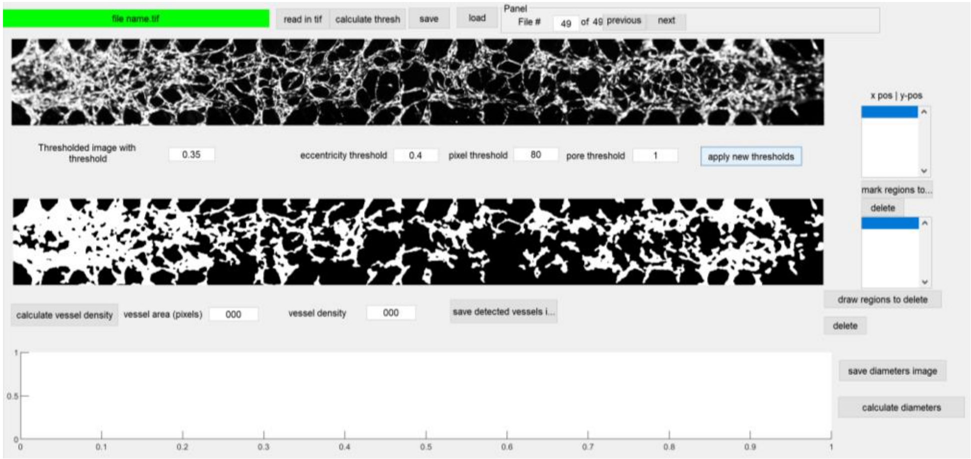

### Supplemental Table 02

| Cell-cell Genes_PC | Cell-cell Genes_FB | Cell-matrix Adhesion_Genes_PC | Cell-matrix Adhesion Genes_FB | Extracellular Matrix Proteins_PC | Extracellular Matrix Proteins_FB |
|---|---|---|---|---|---|
| NLGN2 | CDH6 | ITGA11 | ADAMTS12 | ANOS1 | LTBP1 |
| CD200 | JAM2 | ECM2 | EDA | COL11A2 | CPN2 |
| CDH10 | SIRPA | SORBS1 | ITGB1BP1 | ASPN | LRRC15 |
| ADAM8 | FOXF1 | ITGA3 | NPNT | ECM2 | TIMP4 |
| CTNNA3 | LMO7 | ITGA1 | RHOA | WNT5A | COLEC12 |
| PIP5K1C | CDH11 | MSLN | HPSE | COL4A1 | TNFRSF11B |
| CXADR | NRCAM | HOXD3 | COL5A3 | COL4A2 | NOV |
| PKP3 | BCL2 | ITGB5 | ADAM9 | LAMB1 | KAZALD1 |
| NPHP1 | PKP4 | BCL2L11 | ITGBL1 | LRRTM3 | CCDC80 |
| SRPX2 | SOX9 | L1CAM | EMP2 | MMP14 | EFEMP1 |
| PDLIM5 | COL14A1 | ITGAV | ITGA7 | COL4A5 | ALPL |
| VMP1 | PKP1 | ADAMTS13 | SNED1 | VASN | COL15A1 |
| SRC | CSTA | ITGA4 | NID2 | DGCR6 | CCBE1 |
| RAPGEF1 | NTN1 | CTNNB1 | VCAM1 | TRIL | TNXB |
| CDH24 | CYR61 | EMILIN1 | FN1 | VIT | NPNT |
| KRT18 | STXBP6 | COL3A1 | | LRRC24 | COL14A1 |
| DSP | THY1 | ITGA6 | | LAMA5 | FBN2 |
| JUP | EGFR | TMEM8B | | COL28A1 | ABI3BP |
| CTNNB1 | CNN3 | ITGA8 | | COL27A1 | ADAMTS1 |
| ITGA6 | NECTIN3 | ITGB3 | | ENAM | FGF10 |
| EPCAM | PAK4 | LYVE1 | | EMID1 | HPSE |
| ITGA8 | S100A11 | CTGF | | ADAMTS5 | MATN2 |
| ARVCF | FAT1 | CD63 | | POSTN | VCAN |
| ICAM1 | VCAM1 | FREM1 | | MMP15 | HSPG2 |
| CDH2 | DLG2 | NID1 | | VWA1 | WISP2 |
| SYNJ2BP-COX16 | | THBS3 | | COL1A2 | NAV2 |
| ICAM5 | | | | MMP16 | RELN |
| COL19A1 | | | | THBS1 | TGFBI |
| FNDC3A | | | | ADAMTS10 | COL5A3 |
| CDH3 | | | | LOXL1 | CD180 |
| PKD1L1 | | | | WNT4 | COL6A1 |
| ESAM | | | | THBS2 | COL6A3 |
| CYFIP2 | | | | MMP2 | TIMP3 |
| PDLIM1 | | | | SPON2 | ADAMTS3 |
| LRRC7 | | | | ADAMTS13 | TGFBR3 |
| FBLIM1 | | | | GLG1 | DCN |
| SHC1 | | | | MGP | EGFL6 |
| | | | | COMP | FLRT2 |
| | | | | LTBP3 | FLRT3 |
| | | | | MMP11 | FMOD |
| | | | | EMILIN1 | PRELP |
| | | | | GPC1 | LTBP2 |
| | | | | ADAMTSL2 | FGFR2 |
| | | | | APOE | HNRNPM |
| | | | | COL7A1 | CYR61 |
| | | | | COL3A1 | TNC |
| | | | | LRIG2 | LTBP4 |
| | | | | LUM | TLR3 |
| | | | | COL5A2 | NID2 |
| | | | | COLQ | COL22A1 |
| | | | | CTHRC1 | MMP1 |
| | | | | COCH | MMP3 |
| | | | | PLSCR1 | FBN1 |
| | | | | COL5A1 | MMP12 |
| | | | | GPLD1 | ELN |
| | | | | FBLN2 | TIMP1 |
| | | | | WISP1 | ADAMTSL5 |
| | | | | LINGO2 | ADAMTSL4 |
| | | | | HPSE2 | COL6A6 |
| | | | | SPARC | SERAC1 |
| | | | | MFAP4 | CLU |
| | | | | COL19A1 | MYOC |
| | | | | COL24A1 | TGFB1 |
| | | | | LRIG3 | NDNF |
| | | | | COL25A1 | COL11A1 |
| | | | | CTGF | SCARA3 |
| | | | | RELL2 | PTX3 |
| | | | | COL21A1 | FN1 |
| | | | | AGRN | |
| | | | | COL9A2 | |
| | | | | ADAMTS2 | |
| | | | | COL4A4 | |
| | | | | ADAMTS4 | |
| | | | | MMP7 | |
| | | | | OLFML2A | |
| | | | | RARRES2 | |
| | | | | WNT2 | |
| | | | | OLFML2B | |
| | | | | MMP10 | |
| | | | | CHADL | |
| | | | | ECM1 | |
| | | | | EGFL7 | |
| | | | | NID1 | |
| | | | | WNT6 | |
| | | | | MMP17 | |
| | | | | LAMC2 | |
